# The Immunogenicity Database Collaborative (IDC): A Standardized, Publicly Available Database for Clinical Immunogenicity Observations and Insights

**DOI:** 10.64898/2025.12.08.692993

**Authors:** Sudhanshu Agnihotri, Bruno Gonzalez-Nolasco, Brinda Monian, Sofie Pattijn, Chloe Ackaert, Patrick Wu, Hubert Kettenberger, Sophie Tourdot, Timothy Hickling, Zicheng Hu, Richard E. Higgs, Daniel S. Leventhal

**Affiliations:** Department of Pharmaceutical Sciences, University at Buffalo, The State University of New York, Buffalo, New York, USA; Early Development Services, Lonza Biologics Inc., Cambridge, MA, USA; Generate Biomedicines, Somerville, MA, USA; In Vitro Immunology, RIqvia Laboratories, Gosselies, Belgium; Department of Translational Pharmacokinetics and Pharmacodynamics, Genentech Inc, South San Francisco, CA, USA; Large Molecule Research, Roche Pharma Research and Early Development, Roche Innovation Center Munich, Penzberg, Germany; Pharmacokinetics, Dynamics and Metabolism, Pfizer Inc., Andover, MA, USA; Pharma Research and Early Development, Roche Innovation Centre Welwyn, Roche, Welwyn Garden City, UK; Quasor Ltd, Loughborough, England, UK; Eli Lilly and Company, Indianapolis, IN, USA; Xaira Therapeutics, South San Francisco, CA, USA; Tactyl LLC, St. Petersburg, FL, USA

## Abstract

The incidence and impact of anti-drug antibodies (ADAs) against biotherapeutics remain difficult to predict, limiting efforts to mitigate immunogenicity risk prior to clinical trials. Existing data are fragmented across disparate sources with inconsistent definitions, representing a key barrier to progress in the field. Here, we present the Immunogenicity Database Collaborative (IDC) and its release of the Immunogenicity Database (DB) V1: a structured clinical immunogenicity dataset integrating therapeutic characteristics, sequence, and patient cohort-level data from publicly available sources. The dataset includes 4,146 ADA datapoints, 1,788 cohorts, 727 clinical trials and 218 therapeutics. We highlight trends in ADA incidence, evaluate sources of variability, and identify driving factors of immunogenicity risk. This work provides a foundational resource to standardize and support immunogenicity risk assessment across the industry. It also provides an initial data architecture and invites the research community to contribute towards future expansions of the database into key areas of interest to the field.

## Introduction

The inclusion of biologics into modern medicine has revolutionized the treatment of human disease. However, the therapeutic potential of protein-based drugs can be undermined by unwanted immunogenicity, where the immune system generates anti-drug antibodies (ADAs) against the biologic^1,2^. ADAs can neutralize drug activity^3^, accelerate clearance^4^, or cause hypersensitivity reactions^5^, ultimately compromising safety and efficacy^6–11^. In some cases, immunogenicity detected late in clinical development can halt otherwise promising therapeutics^12^. These risks underscore the importance of deepening our understanding of the root causes of immunogenicity, as well as continually advancing preclinical risk assessment tools and mitigation strategies.

Immunogenicity risk is influenced by multiple factors that can be broadly categorized into product-, treatment-, and patient-related variables^13,14^. Many product-intrinsic risks can be assessed preclinically, including a drug’s mechanism of action (MOA), antigenicity (e.g., T and B cell epitope content), developability attributes (e.g., aggregation propensity, polyreactivity, etc.), and drug product critical quality attributes (CQAs) (e.g., impurity content, host-cell protein levels, etc.)^15^. Treatment-related factors include a drug’s dose quantity, dose frequency and route of administration (ROA), while patient-related factors include immune status, concomitant medications, and genetic background^16^. The complex interplay between these variables can further impact ADA outcomes.

While these factors are well established as independent contributors, their complex, context-dependent interplay with ADA outcomes makes it challenging to deconvolute their relative contributions to immunogenicity risk. For example, a non-human, highly antigenic protein may not trigger ADA formation in an immunosuppressed population, while a fully humanized therapeutic may elicit strong responses if its target is expressed on antigen-presenting cells^16,17^. A wide range of clinical and ADA assay technical factors can influence reported immunogenicity outcomes, therefore regulatory guidance has historically discouraged direct comparison of ADA rates across therapeutics^18^. Nonetheless, when scale meets sufficient methodological and clinical context, such analyses can yield valuable scientific insights, and thus high-quality, fit-for-purpose datasets that link immunogenicity outcomes to relevant risk factors are of great value to the field.

Prior publicly available clinical immunogenicity data are fragmented across FDA labels, ClinicalTrials.gov entries, EMA reports, and throughout primary literature. It often contains inconsistent reporting of ADA outcomes (e.g., transient ADA, treatment-emergent ADA, etc.) and limited methodological details for ADA assessments. Published aggregated datasets utilized in the field frequently report a single ADA frequency and often differing values for the same molecules (Fig S1)^19–22^. This heterogeneity impairs data accessibility, hinders context aware comparisons and limits the ability to elucidate drivers of immunogenicity risk.

To address these gaps, we established the Immunogenicity Database Collaborative (IDC) and present its first public release: a curated, harmonized, and publicly accessible clinical immunogenicity database (the IDC DB V1). Here, we describe the dataset architecture, characterize its contents, and demonstrate its utility by exploring variable contributions to ADA frequency across biologics and a broad range of clinical contexts.

## Results

### Creating a fit-for-purpose data architecture

The IDC database was built using TIDY data principles^23^ and refined through multiple rounds of pilot testing and expert feedback. The architecture consists of three relational data tables: Therapeutics, Sequences, and Clinical Trials.

**The Therapeutics table** (Table S1) assigns a unique Therapeutic ID to each drug and captures a range of high-level attributes spanning molecule features, targets, mechanism of action, regulatory status, and biosimilar-to-originator relationships. Information was aggregated from various sources including FDA and EMA product labels, WHO INN databases^24^, and curated drug repositories such as DrugBank^25^ and AdisInsight^26^. The IDC DB V1 reflects information up to the end of 2024.

**The Sequences table** (Table S2) provides amino acid sequences organized by chain or domain, providing a unique Sequence ID and linking to a parental Therapeutic ID. Sequence verification was attempted by confirming identity in at least two independent sources, including IMGT^27^, KEGG^28^, Drugs@NCATS^29^, Thera-SabDab^30^, or directly in patents. When sequences were not available, entries were flagged accordingly.

**The Clinical Trials table** (Table S3) captures trial- and cohort-level features with respective unique identifiers. It catalogs treatment regimens, patient cohort characteristics, assay properties, ADA/neutralizing ADA (nADA) frequencies, assessment timepoints, and a variety of other features and metrics. Reported ADA measures were unified into an ADA frequency at the assessment timepoint (see Supplementary Information for further details). While ADA incidence requires individual-level longitudinal data, this dataset uses ADA frequency as a practical alternative to the available summary-level data most commonly reported in ClinicalTrials.gov, FDA labels, and the EU Clinical Trials Register.

Together, these components establish IDC DB V1 (Table S4), a framework for standardized immunogenicity data capture and enable side-by-side exploration of clinical, mechanistic, and molecular features.

### Clinical immunogenicity data set overview

The IDC DB V1 features 218 therapeutics (e.g., drug products) with 142 for which clinical trial data was available (Fig. 1a). In total, the database integrates 1,788 cohorts across 727 clinical trials, yielding 4,146 ADA-related datapoints: 3,334 ADA frequency, 621 nADA frequency, 147 ADA titer, and 44 nADA titer measurements. This scale enables comparative analyses across therapeutic classes and clinical contexts.

**Figure 1:**
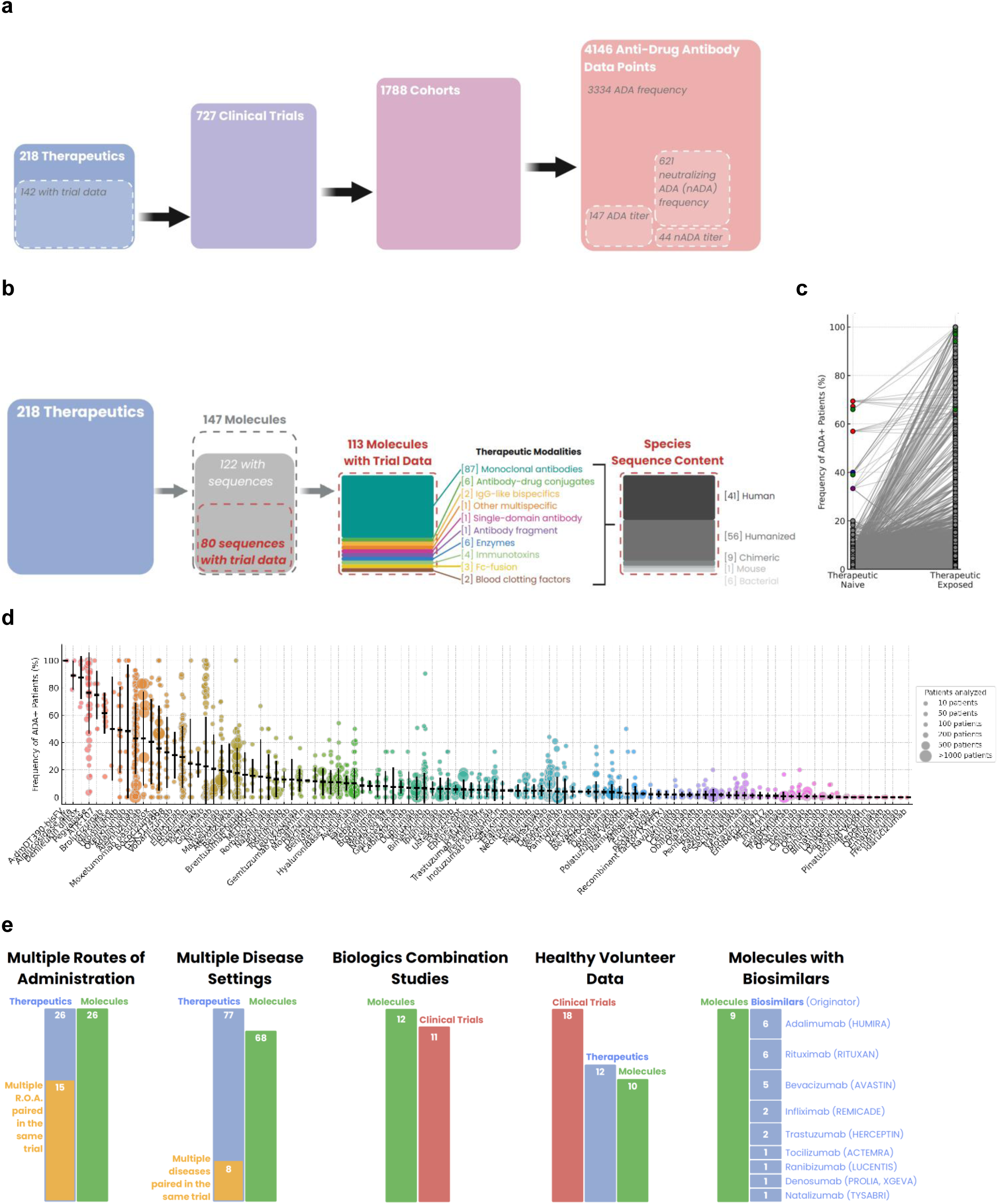
An overview of the clinical immunogenicity database. (a) The dataset features 218 therapeutics with 142 having clinical trial data. In total, the database integrates 1,788 cohorts across 727 clinical trials, yielding 4,146 ADA-related datapoints: 3,334 ADA frequency, 621 nADA frequency, 147 ADA titer, and 44 nADA titer measurements. (b) Of the 218 therapeutics captured, there are 148 unique molecules with 122 sequences identified for those molecules. Eighty sequences have clinical trial data captured. A variety of protein modalities and source species are represented amongst the 113 molecules having clinical trial data captured, the vast majority representing monoclonal antibodies. (c-d) Reported ADA+ patient frequency for each datapoint is shown. (c) Datapoints are separated based on labeling as coming from patients not exposed to a therapeutic (Therapeutic Naive) or those exposed to a therapeutic (Therapeutic Exposed). Red = Brolucizumab; Blue = Otelixizumab; Purple = Utomilumab, Green = Denileukin Difitox. (d) Datapoints are grouped and colored by therapeutic. The size of each datapoint is proportional to the number of patients assessed for ADAs within the patient cohort at that specific timepoint. Mean ADA+ frequency and standard deviation are shown. (e) The number of therapeutics, molecules and clinical trials captured within the dataset are shown for each of the given context.

The composition of molecules (e.g., unique proteins) captured in the dataset is summarized in Fig. 1b. Monoclonal antibodies predominate (n=87), with several antibody–drug conjugates (n=6), bispecific or multispecific antibodies (n=3 total), single-domain/fragment antibodies (n=2), enzymes (n=6), immunotoxins (n=4), Fc-fusion proteins (n=3), and blood factors (2n=). Of the 147 molecules captured, 122 had amino-acid sequences available and 80 of those have clinical trial data captured in the dataset. The species origin for molecules captured skews toward humanized (n=56) and human (n=41), with smaller counts of chimeric (n=9), mouse (n=1), and bacterially derived proteins (n=6).

Immunogenicity datapoints were labeled as originating from either therapeutic-naïve or therapeutic-exposed cohorts. While most ADA frequencies increase at timepoints post exposure, several molecules (Brolucizumab, Otelixizumab, Utomilumab, and Denileukin Diftitox) exhibited elevated pre-existing ADA in naïve patients at baseline (Fig. 1c). Unless otherwise noted, all subsequent analyses were focused on therapeutic exposed populations only.

The therapeutics captured in the Clinical Trials table cover a wide range of ADA frequencies and illustrate the variability of ADA frequencies reported for the same therapeutic across trials, cohorts and timepoints (Fig. 1d). Fluctuations in reported ADA rates may reflect multiple contributing factors, thus these findings underscore the importance of interpreting immunogenicity data within the specific clinical context and conditions under which data were obtained. To track immunogenicity rates at multiple levels, we also provide aggregated ADA frequencies by cohort (Fig. S2a), trial (Fig. S2b), therapeutic (Fig. S2c), and molecule (Fig. S2d), with data provided in Table S5.

Finally, the data set provides opportunities to examine key features of great interest to the field (Fig. 2e). Examples include 26 therapeutics with multiple routes of administration (ROA) (15 trials with multiple ROA in the same trial), 77 used within multiple indications (8 within the same trial), 11 used in combination, 18 studies in healthy volunteers, and 9 molecules with biosimilars. These groupings define the analytic space for some representative evaluations presented in the next section and highlight opportunities for the immunogenicity community to further expand the dataset.

**Figure 2:**
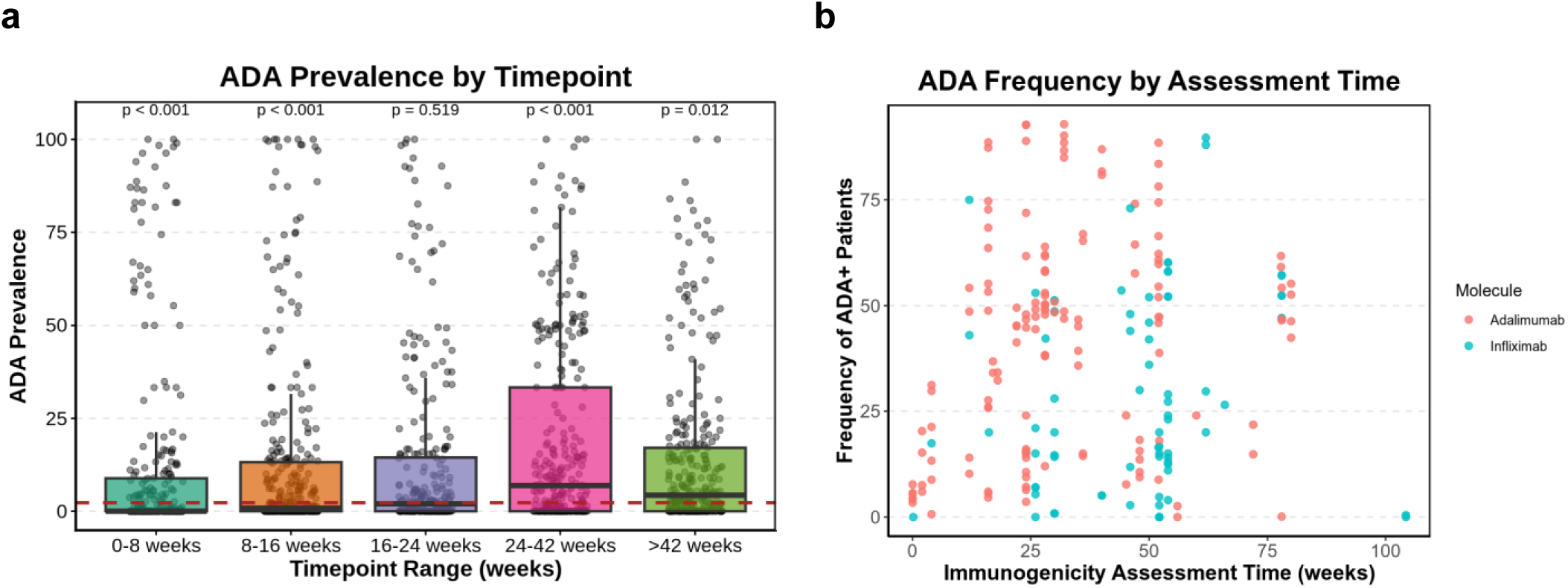
ADA frequency measurements over time. (a) Distribution of ADA frequencies across clinical cohorts plotted by weeks since first therapeutic exposure. Each point represents a cohort-level measurement at a reported timepoint. 1000-fold bootstrapping was used to test if the median of each group is significantly different from the overall median. (b) Examples from Adalimumab and Infliximab clinical trials showing ADA frequencies measured at multiple timepoints, with values plotted relative to weeks post first dose.

#### Insights into drivers of immunogenicity risk

Substantial variability in anti-drug antibody (ADA) frequency was observed across clinical cohorts in the IDC dataset, highlighting both inter- and intra-therapeutic heterogeneity (Fig. S2a). Some therapeutics demonstrated consistently high ADA frequencies across trials (e.g., Denileukin Diftitox), while others exhibited broad variability, potentially reflecting trial- and cohort-specific factors (e.g., Adalimumab, Infliximab). At scale, these differences raise fundamental questions: What drives ADA frequency variation across biologics? Which drug-intrinsic and drug-extrinsic features matter most? To begin addressing these, we evaluated the impact of several hypothesized drivers of clinical immunogenicity using both univariate patterns and case-specific examples.

ADA sampling schedules captured in the database exhibit an expectedly wide range, with some cohorts assessed within a week of dosing and others up to several years later. ADA frequency generally increased over time, peaking around 24-42 weeks after the first dose (Fig. 2a). Early elevated ADA frequencies may reflect prior exposure or in rare cases cross-reactive pre-existing ADAs (Fig. 1c). Analyses of Adalimumab and Infliximab (Fig. 2b) further illustrate this trend, with both therapeutics demonstrating rising ADA frequencies in later post-dose samples and peaking at 25-50 weeks. These observations reinforce the impact sampling time has on the reported frequency of ADA within a cohort. For this reason, we generated additional data tables listing the maximum ADA frequency observed for a cohort within a trial (cohort level), the average ADA frequency of all cohorts within a trial (trial level) and the average ADA frequencies calculated by aggregating all ADA positive and negative patients amongst all cohorts for a particular drug (therapeutic level) or protein (molecule level) (Table S5).

#### Immunogenicity and the Role of Therapeutic Context

The clinical setting in which a therapeutic is delivered (e.g. the dose amount, dosing interval and ROA) can potentially impact ADA induction rates. Across the dataset, higher ADA frequencies were observed at lower dose levels, with the highest seen for >0-60 mg per dose (Fig. 3a). These findings may reflect lower immune tolerance thresholds at subtherapeutic exposure or potential assay interference due to higher drug levels at sampling. However, when evaluating a smaller set of therapeutics all of which target the PD-1/PD-L1 pathway in oncology settings (Durvalumab, Atezolizumab, Avelumab, Pembrolizumab), higher ADA frequencies were observed at intermediate dosing levels (700-1200 mg) (Fig. 3b). Previous reviews evaluating the immunogenicity of these kinds of immunotherapics had not determined a dose-dependent effect on nADA incidence, however this was likely due to limited available data^31^.

**Figure 3:**
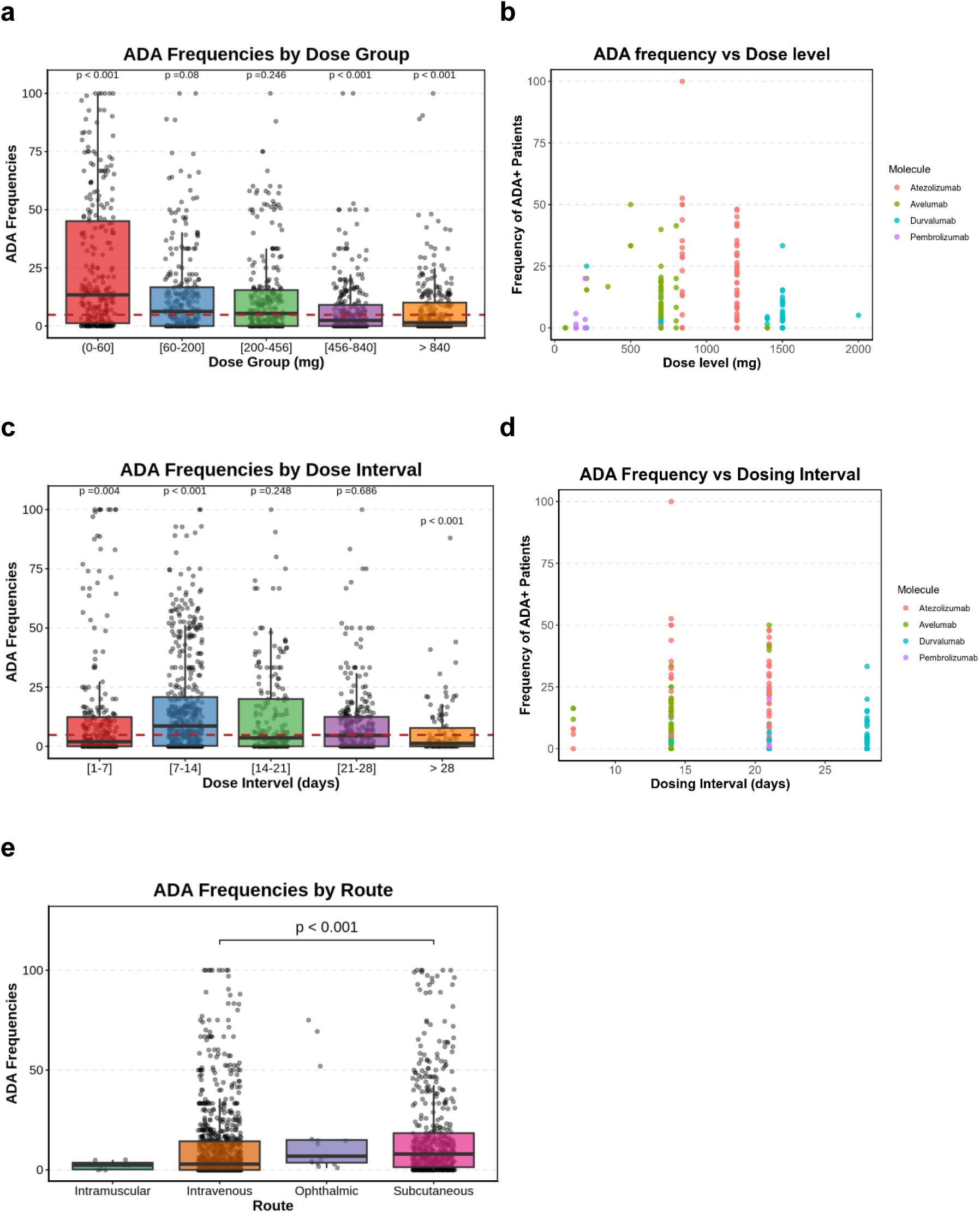
Impacts of dose amount, dose interval and route of administration on ADA frequency. (a) ADA frequencies across cohorts grouped by dose amount categories. Each point represents a cohort-level measurement at a reported timepoint. A 1000-fold bootstrapping approach was used to test whether the median of each group differs significantly from the overall median. (b) ADA frequencies for Durvalumab, Atezolizumab, Avelumab, and Pembrolizumab cohorts, stratified by dose amount. Each point represents a cohort-level measurement at a reported timepoint. (c) ADA frequencies across cohorts grouped by dosing interval categories. Each point represents a cohort-level measurement at a reported timepoint. A 1000-fold bootstrapping approach was used to test whether the median of each group differs significantly from the overall median. (d) ADA frequencies for Durvalumab, Atezolizumab, Avelumab, and Pembrolizumab cohorts, stratified by dosing interval. Each point represents a cohort-level measurement at a reported timepoint. (e) ADA frequencies across cohorts stratified by route of administration (subcutaneous vs. intravenous). Each point represents a cohort-level measurement at a reported timepoint. A 1000-fold bootstrapping approach was used to test whether the median differs significantly between subcutaneous and intravenous routes.

Regarding dose interval, we observed a peak in ADA frequency for regimens administered at approximately 7-14 day intervals (Fig. 3c), a pattern reminiscent of standard prime-boost immunization schedules. A similar trend was observed for the smaller set of anti-PD-1/PD-L1 therapeutics at 14- and 21-day intervals (Fig. 3d), suggesting that immune system re-exposure timing may modulate the likelihood of ADA formation. The ROA may also impact immunogenicity, with subcutaneous (SC) administration classically thought to be more immunogenic than intravenous (IV) due to enhanced uptake by antigen-presenting cells in peripheral tissues^14^. In the IDC DB V1 dataset, SC routes were associated with modestly higher ADA frequencies than IV (Fig. 3e). There are currently only 3 intramuscular and 4 ophthalmic administered therapeutics in the IDC DB V1, limiting the insights which can be gained for those ROA. Only a small subset of therapeutics had datapoints associated with both SC and IV administration, with an even smaller subset providing comparative data within the same clinical trial (Fig. S3). Higher ADA frequencies were observed for SC administered therapeutics, especially those with higher overall ADA frequencies (e.g. Adalimumab, Bococizumab and Golimumab); however, those evaluated within the same trial exhibited minimal differences between ROA. Together, these findings suggest that ROA may represent a contributing factor to ADA risk, however, additional within-trial comparisons are required to conduct more robust evaluations and draw definitive conclusions.

#### Patient Immune Status is Key

A biologic’s mechanism of action (MOA) can influence its immunogenicity both directly via immune system modulation, and indirectly based on the pre-existing immune status of the patient population in which it is used^21^. When grouped by immunomodulatory profile, biologics with T and B cell activating MOAs (ex. checkpoint inhibitors) had higher average ADA frequencies than those with T and B cell depleting or immunologically neutral MOAs (Fig. 4a). While therapeutics having anti-inflammatory MOAs had the highest overall ADA frequencies (Fig. 4a), this may be confounded by the heightened immune status of the patient population or increased antigen presenting cell uptake due to expression of the target receptor. Indeed, patients with inflammatory or autoimmune conditions exhibited the highest ADA frequencies compared to other disease indication cohorts (Fig. 4b).

**Figure 4:**
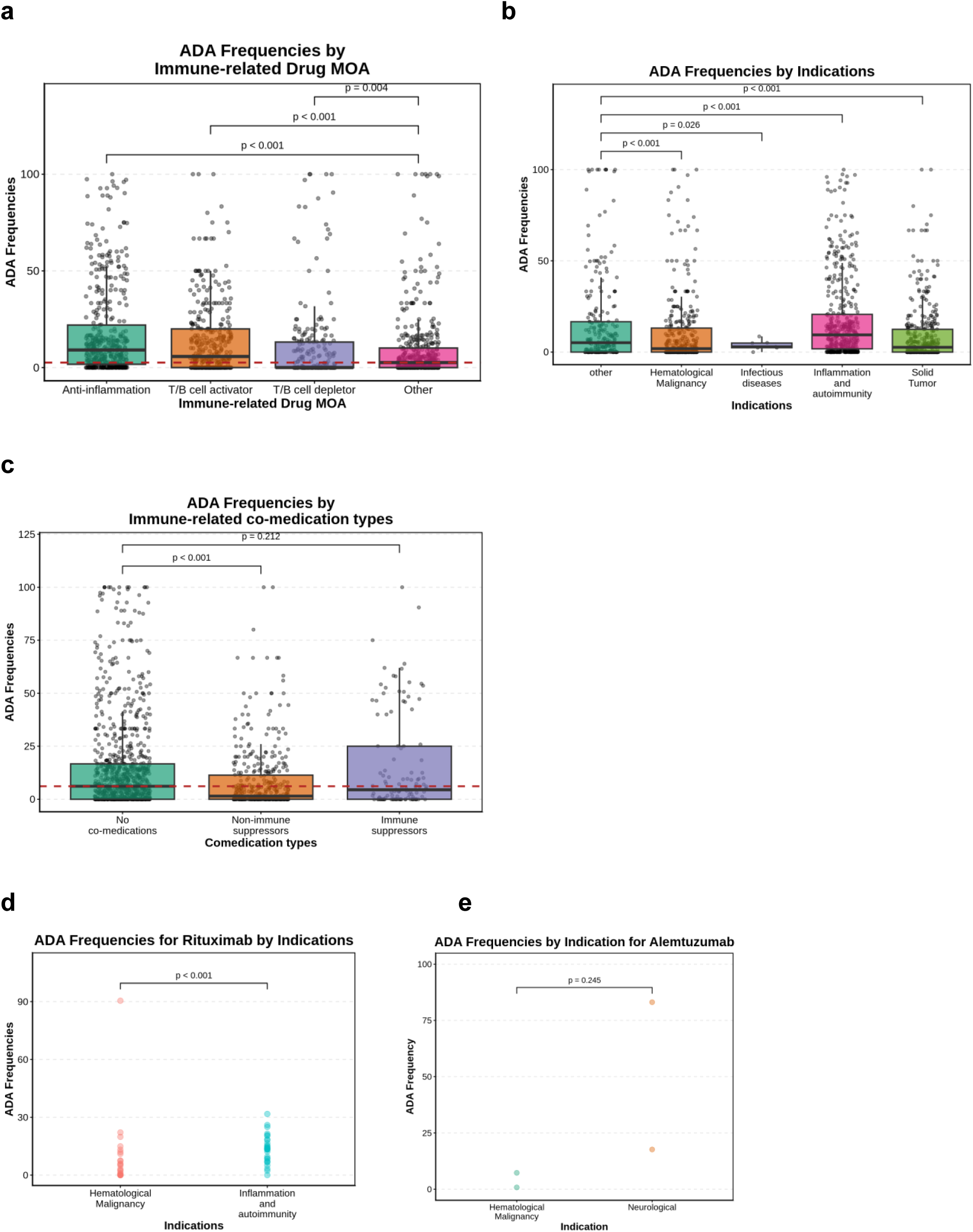
ADA frequencies observed across various drug and comedication mechanisms of action and disease indications. (a) ADA frequencies across biologics grouped by immune mechanism of action. Each point represents a cohort-level measurement at a reported timepoint. The “other” group was used as the baseline, and a 1000-fold bootstrapping approach was applied to test whether the median of each group differs significantly from the baseline. (b) ADA frequencies across patient cohorts grouped by disease indication. Each point represents a cohort-level measurement at a reported timepoint. The “other” group was used as the baseline, and a 1000-fold bootstrapping approach was applied to test whether the median of each group differs significantly from the baseline. (c) ADA frequencies across cohorts stratified by comedication type. Each point represents a cohort-level measurement at a reported timepoint. The “no-comedications” group was used as the baseline, and a 1000-fold bootstrapping approach was applied to test whether the median of each group differs significantly from the baseline. (d–e) ADA frequencies for Rituximab (d) and Alemtuzumab (e) cohorts stratified by disease indication. Each point represents a cohort-level measurement at a reported timepoint. Wilcox tests were used to calculate the p values in d and e.

Concomitant medications frequently differ between patient populations and can significantly impact immune status and ADA responses. When evaluated in aggregate, cohorts labeled as having received immunosuppressive agents (e.g., methotrexate or corticosteroids) or no comedications exhibited higher average ADA frequencies than cohorts treated with non-immune suppressive co-medications (Fig. 4c). While this finding contradict previous reports^32^, these co-medications are often utilized in autoimmune patients who exhibit increased ADA frequencies (Fig. 4b). When evaluated on a per therapeutic basis the inclusion of suppressive co-medications most often (4 of 5 therapeutics) led to reduced ADA frequencies (Fig. S4). However, no apparent differences were observed for the very few datapoints with cohorts evaluated within the same trial and thus inclusion of additional intratrial evaluations of co-medications remain a priority for future iterations of the database.

To further evaluate the impact of patient cohort immune status on ADA frequency, we compared two molecules used in different disease settings: Rituximab (an anti-CD20 murine-human chimeric monoclonal antibody that depletes B cells) and Alemtuzumab (an anti-CD52 humanized monoclonal antibody that depletes T and B cells). In the dataset these biologics exhibited higher ADA rates in autoimmune compared to oncology patient cohorts (Figs. 4d–e). This observation further emphasizes the need to interpret immunogenicity endpoints in the context of all relevant clinical features.

#### T cell Epitope Content and Sequence Based Risks

Intrinsic sequence features also represent a major risk factor for developing unwanted immunogenicity. One such feature involves the quantity and quality of T cell epitopes contained within the biologic’s amino acid sequence. Using an industry standard peptide presentation model (NetMHCIIpan-4.X^33^), we enumerated the total number of predicted MHC class II presented peptides containing non-germline encoded residues (e.g. predicted CD4+ T cell epitope load) for each therapeutic (see Methods) and compared it to cohort-level ADA frequencies (Fig. 5a). Higher predicted CD4+ T cell epitope counts were weakly but positively correlated with ADA rates, consistent with prior reports utilizing a similar methodology^34^. These results suggest that epitope-based modeling can provide useful predictions of clinical immunogenicity but provide an incomplete perspective when evaluated in isolation. Additional variables such as differences in protein structure, post-translational modifications, host cell protein content^35^, and other drug product-related features may also modulate the immunogenic response, however these variables fall outside the scope of the current dataset.

**Figure 5:**
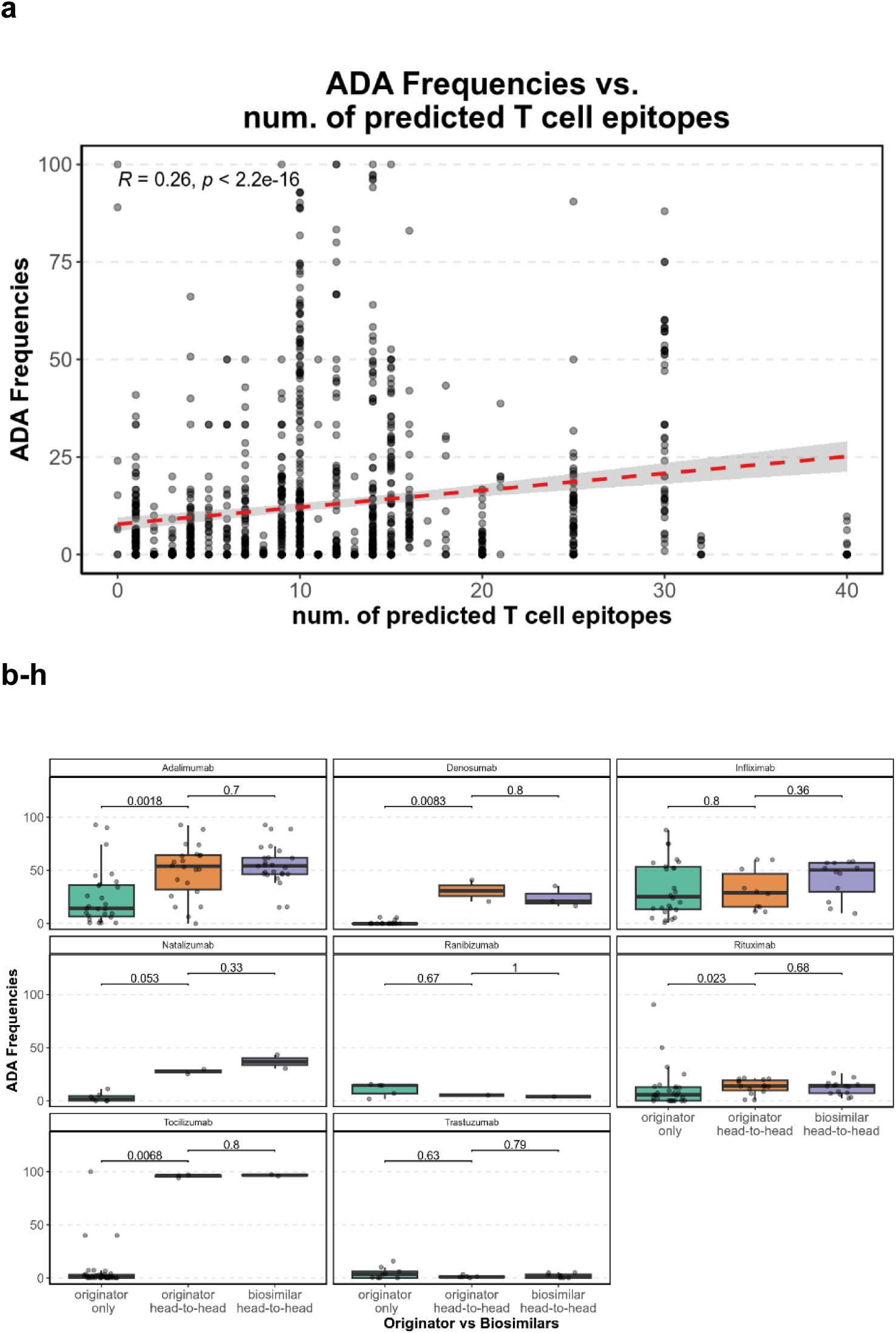
Correlation of T cell epitope content on ADA frequencies and comparison of sequence identical biosimilars. (a) Relationship between predicted CD4⁺ T cell epitope counts and ADA frequencies across therapeutics. Each point represents a therapeutic-level measurement. Linear regression was used for testing the association. (b–h) ADA frequencies reported for an originator drug evaluated in a trial on its own (originator only) and originator–biosimilar pairs evaluated in the same clinical trial (head-to-head). Each point represents a cohort-level measurement at a reported timepoint. t tests were used to compare ADA frequencies between originators and biosimilars.

Biosimilars share the same primary amino acid sequence with the originator or reference product and are required to demonstrate no clinically meaningful differences in safety, purity, or potency^36^. Any differences in ADA frequency observed between biosimilars and reference products would thus likely stem from minor variations in CQAs, process-related impurities, formulation or manufacturing differences, bioanalytical assay design, or trial context. Across seven originator–biosimilar pairs in the IDC DB V1, ADA frequencies were generally comparable when both products were evaluated within the same trial (Fig. 5b-h). We also observe that originators typically reported lower ADA frequencies during their initial clinical development, which may reflect historical assay formats^37^ or differences in patient selection. These observations are in line with regulatory trends towards reducing potentially unnecessary clinical measures during evaluation of biosimilar products with historically minimal immunogenicity liabilities^38^ and underscore the value of organizations publicly reporting immunogenicity metrics.

#### Quantifying Variable Contribution Through Multivariate Analysis

To assess the relative contribution of each variable while accounting for confounders, we performed multivariate regression using a range of clinical and therapeutic features captured in the dataset. As shown in Fig. 6 and Table S6, the top three variables associated with higher ADA frequency were therapeutic MOA, disease indication and predicted T cell epitope content. Conversely, variables such as ROA, co-medication immunomodulation type and dose level had weaker or inconsistent associations similar to the year in which the trial was completed. These analyses highlight the multifactorial nature of immunogenicity and support current integrative risk assessment practices that combine clinical, mechanistic, and sequence-based inputs.

**Figure 6:**
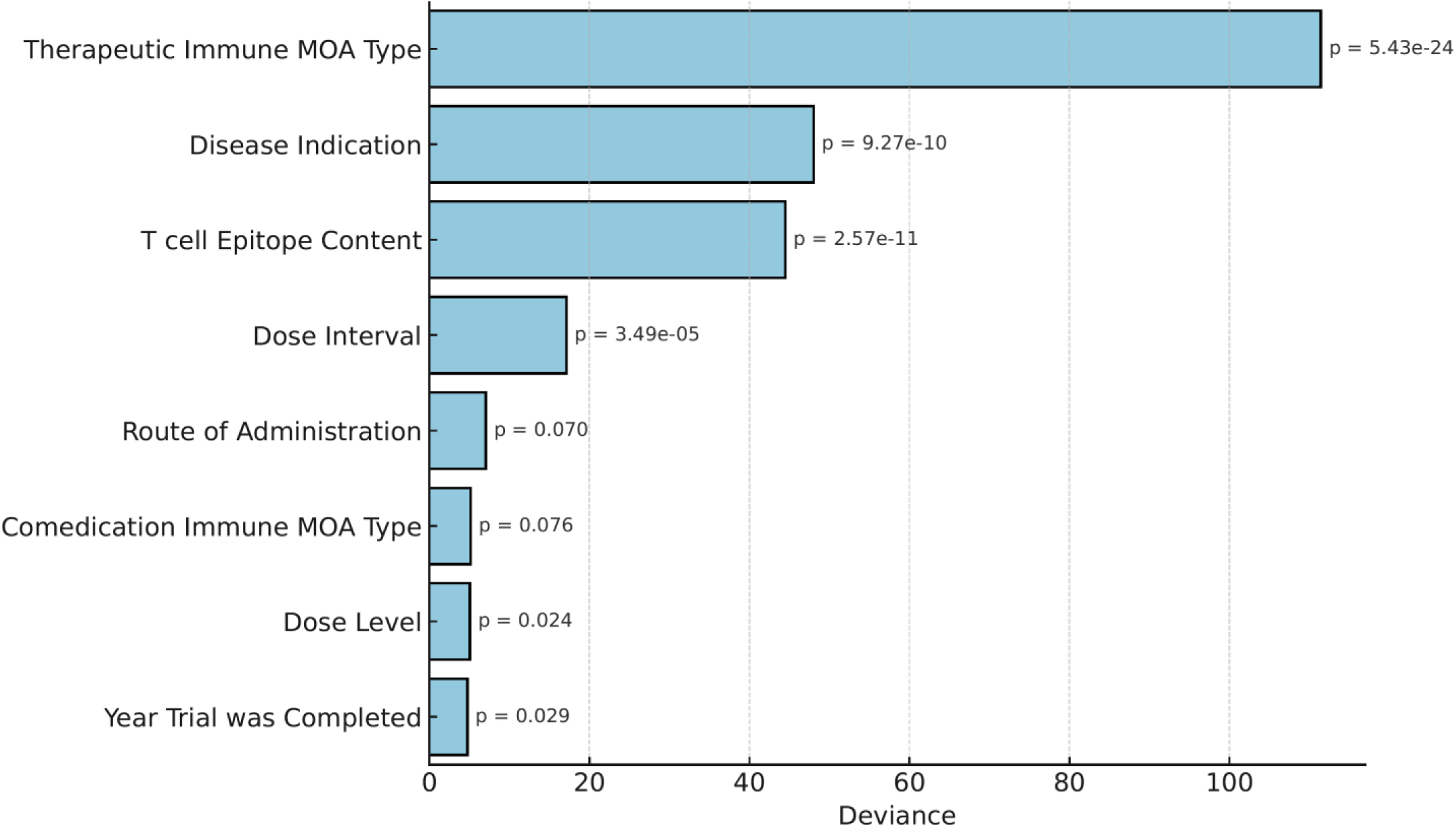
Multivariate analysis for drivers of immunogenicity risk. Multivariate logistic regression of cohort-level ADA frequencies dichotomized into low (<10%) and high (≥10%) categories. Variables included drug mechanism of action, disease indication, predicted T cell epitope content, dosing interval, route of administration, comedication type, dose level, and trial year. The relative importance of each factor was quantified by its contribution to the reduction of model deviance.

## Discussion

The IDC was established to provide a standardized, publicly accessible dataset linking therapeutic, sequence, and clinical features to clinical immunogenicity outcomes. This first release integrates over one hundred therapeutics and thousands of cohorts, capturing ADA frequencies at multiple levels of aggregation. The dataset offers flexible architecture for interoperability and future expansion. Its scale and diversity enable systematic evaluation of immunogenicity risk factors, while also underscoring the multifactorial and context-dependent nature of ADA development.

Insights from the IDC DB V1 both confirm and extend prior observations in the field. SC administration is often considered more immunogenic than IV^39^. Across the dataset, SC delivery was associated with modestly higher ADA rates (Fig. 3e), but significant differences were observed for only 4 of the 24 therapeutics having data from both ROA. This pattern mirrors Davis et al., who reported increased risk for 6 of 20 therapeutics, and Felderman et al., who found no significant correlation when controlling for confounders^40,41^. These findings indicate that ROA may contribute to immunogenicity but is rarely a dominant driver, and more paired SC/IV data, ideally from the same trials, are needed for robust conclusions.

Patient immune context could influence ADA outcomes. Therapeutic MOA, disease indication, and co-medications often overlap, complicating attribution of individual effects. Immunosuppressive concomitant medications are likely to influence ADA frequencies, however, outcomes in our dataset were variable as trials in this dataset were skewed towards autoimmune and oncological indications. Biologics with anti-inflammatory MOAs are predominantly tested in autoimmune settings, where underlying immune status differs substantially from that of oncology cohorts. Case studies such as Rituximab and Alemtuzumab illustrate this point (Fig. 4d and 4e), underscoring the importance of contextual interpretation when considering immunogenicity risk across different indications and regimens.

Sequence features remain key determinants of immunogenicity. Consistent with similar, recent analyses^34^, predicted CD4+ T cell epitope load showed a weak but significant correlation with ADA frequency via univariate analyses (Fig. 5a) and was highlighted as a major contributor via multivariate analyses (Fig. 6). Biosimilars, which are expected to share the same primary amino acid sequence as their reference products, exhibited comparable ADA rates when tested within the same trials, demonstrating a high degree of consistency when a significant subset of factors is controlled for (Fig. 5b-h). Differences between early originator trials and later biosimilar studies likely reflect historical variation in assay format, sensitivity, cut-off determination and patient selection. Greater availability of detailed assay characteristics would substantially improve understanding across the field.

Multivariate regression highlighted four leading variables associated with ADA risk: therapeutic MOA, disease indication, predicted epitope content, and dosing interval (Fig. 6). The top three were also identified in the same order by Zicheng et al. in a smaller but more controlled dataset, providing independent validation^34^. In contrast, route of administration, comedication type, and dose level showed weaker or inconsistent associations.

Several limitations of the IDC DB V1 warrant acknowledgement. ADA outcomes were harmonized to a single baseline or treatment-emergent frequency per cohort-timepoint combination (see Methods), enabling comparability but excluding alternative measures such as transient, persistent or total ADA incidence. Single, aggregate therapeutic- or molecule-level values are provided for convenience (Table S5) but should not be used in isolation, as they obscure heterogeneity arising from trial design and cohort context. ADA assay parameters (including sensitivity, drug tolerance, and antigen source in the case of biosimilar trials) were infrequently reported, limiting the depth of interpretation and the ability to assess the quality of the ADA data reported. Reported ADA frequencies can evolve as clinical development progresses, with larger, more diverse populations and updated assay technologies often providing higher sensitivity and drug tolerance. In the absence of assay drug tolerance and pharmacokinetic measures, it is not possible to exclude potential impacts of drug exposure (e.g. serum drug concentrations) on reported ADA measures. Biosimilarity and interchangeability studies can also report frequencies that may differ from early trials. Thus, utilization of trial start and end dates can provide longitudinal perspective on reported ADA frequencies. Finally, nADA frequencies, ADA titers, and clinical outcomes such as efficacy or safety impacts hold substantial value for understanding immunogenicity risk, however these datapoints were infrequently reported, minimizing meaningful conclusions that can be drawn beyond total ADA frequency.

Despite these caveats, the IDC DB V1 represents a significant advancement over prior published datasets^19–22^, integrating a broad set of molecules, clinical contexts, and mechanistic features into a unified framework. The database supports both large-scale comparative analyses and targeted case studies, facilitating identification of consistent immunogenicity drivers while also illustrating the limitations of single-factor interpretations.

Future expansions will extend the database across several dimensions. First, broader clinical coverage, including additional biosimilars, non-antibody modalities (e.g., peptides, Fc-fusions, nanobodies, enzymes), inclusion of anonymized individual patient data and features such as multiple ROA, immunomodulatory concomitant medications and measures of impact on clinical outcomes. Second, more detailed assay metadata, particularly sensitivity and drug tolerance, to strengthen cross-trial observations. Third, incorporation of non-clinical risk features such as biophysical properties (aggregation propensity, polyspecificity, post-translational modifications) and preclinical risk assessment readouts (dendritic cell uptake, MHC II-associated peptide proteomics, T cell antigenicity, etc.). Challenges in harmonization of datasets will remain, but industry-wide collaborative efforts offer a path forward with approaches such as federated learning providing an avenue to leverage sensitive datasets without compromising data privacy^42^.

The release of the IDC DB V1 showcases both the current state of the database architecture and the constraints of available public data. Ongoing community engagement will be essential to guide expansion, refinement, and prioritization of new features. By contributing data and feedback, the field can ensure that IDC DB evolves into a durable shared resource for identifying predictors of immunogenicity, benchmarking risk assessment methods, and improving the safe and effective use of biologics.

## Methods

### IDC Sources

The IDC was constructed through domain expert-led systematic extraction and curation of publicly available information on clinical immunogenicity. Primary data sources included U.S. Food and Drug Administration (FDA) and European Medicines Agency (EMA) product labels, ClinicalTrials.gov, the EU Clinical Trials Register, WHO International Nonproprietary Name (INN) databases, and curated repositories such as DrugBank^25^ and Thera-SAbDab^30^, supplemented by peer-reviewed publications, patent filings, and regulatory submissions where available. Data were organized into three interlinked tables, Therapeutics, Sequences, and Clinical Trials, each connected by unique identifiers to enable relational mapping.

### Sections of IDC DB V1

The Therapeutics table (Table S1) captured drug-level attributes, including INN, trade names, antibody backbone (e.g., IgG1, IgG4, Fc-fusion), species origin (murine, chimeric, humanized, or human), expression system, disease indication, molecular target, and mechanism of action.

Mechanistic characteristics were recorded to highlight drugs that are inherently immunogenic, such as immune system activators that confer heightened risk in certain patient populations. Regulatory milestones were also documented, including FDA and EMA approval status and timelines when obtainable. Each therapeutic was assigned a unique Therapeutic_ID that served as the primary key across tables. Biosimilars were annotated in the field “Labelled as Biosimilar?” and linked to their parent molecule through the “Progeny of” field, while sequence cross-references were captured through Sequence IDC Identifiers.

The Sequences table (Table S2) recorded amino acid sequences for therapeutic agents, organized by heavy and light chains and linked to their parent therapeutic via Therapeutic_ID. Each sequence was annotated with chain type, sequence source, and domain identity, and curated primarily from IMGT^27^, regulatory filings, and patent databases, with additional references from repositories such as the RCSB Protein Data Bank^43^. For biosimilars, only the parent drug sequence was reported since by definition the primary sequences should be identical. A limitation of this table is that sequences for certain novel constructs or investigational biologics were not publicly available.

The Clinical Trials table (Table S3) formed the core of the dataset, capturing immunogenicity outcomes at the cohort level within each trial. Recorded variables included trial identifiers (e.g., NCT or EudraCT numbers), drug regimen, dosing, route of administration, patient population characteristics, therapeutic exposure status (treatment-naïve versus treatment-exposed), ADA frequency, neutralizing antibody data, timepoints of ADA assessment, and assay characteristics such as format, sensitivity, and drug tolerance when reported.

One of the primary challenges in curating the IDC DB V1 was the varied measurements and interpretations used to report ADA outcomes. Terms such as total, treatment-emergent, transient, persistent, and treatment-boosted ADA were reported inconsistently across sources. To harmonize this, only two frequency types were retained or derived where possible: (1) treatment-naïve (i.e., all patients ADA-positive in unexposed cohorts) or baseline (i.e., all patients ADA-positive at baseline), and (2) treatment-emergent (including both treatment-induced and treatment-boosted rates).

The term “ADA frequency” is used instead of “ADA incidence” in this database to denote the percentage of patients who tested positive for ADAs at a given timepoint. ADA incidence, as defined by Shankar et al. (2014), refers to the number of patients who developed treatment-emergent ADA (that is, treatment-induced or treatment-boosted) during a study^44^. Providing an accurate measure for incidence requires individual patient-level information, including baseline and follow-up ADA status. However, most data sources provided only summary-level data in the form of ADA positivity at discrete time points without individual patient profiles or baseline status. This limitation necessitated the use of ADA frequency as a more general and descriptive metric to the proportion of ADA-positive patients. ADA frequency is then interpreted in the context of therapeutic exposure. In therapeutic-naïve patients, frequency denotes any ADA positivity at baseline or in placebo or comparator cohorts at a given timepoint. In therapeutic-exposed patients, ADA frequency is interpreted as treatment-emergent ADA at a given timepoint.

### T cell epitope prediction

Potential immunogenic CD4⁺ T cell epitopes were predicted computationally using NetMHCIIpan-4.3 EL^45^ for nine common HLA-DR alleles: HLA-DRB1*01:01, HLA-DRB1*03:01, HLA-DRB1*04:01, HLA-DRB1*07:01, HLA-DRB1*08:01, HLA-DRB1*09:01, HLA-DRB1*11:01, HLA-DRB1*13:01, and HLA-

DRB1*15:01. These alleles were selected based on their broad global population coverage (85.61%) and the availability of extensive MAPPs training data ^33,46,47^. Peptides ranging from 12 to 20 amino acids were considered to be strongly presented on HLA-DR if the predicted binding core (a nine-mer peptide) had a percentile rank score within the top 10%. To filter likely tolerized epitopes, the predicted binding cores were queried against the natural antibody repertoire in the Observed Antibody Space (OAS) using the BioPhi humanness analysis tool ^48,49^. Binding cores found in more than 110 human OAS subjects were excluded from further analysis. Binding cores were also removed if their sequences matched nine-mer peptides from the human reference proteome^50^. In addition, cores overlapping with ‘knobs-into-holes’ mutations were excluded, as such mutations have been suggested to have a lower risk of immunogenicity^51,52^. The remaining binding cores were retained as potentially immunogenic CD4⁺ T cell epitopes.

### Statistical Analysis

The association between anti-drug antibody (ADA) frequency and individual immunogenicity risk factors was investigated. Initial analysis revealed that ADA frequencies significantly deviated from a normal distribution. Consequently, we employed a non-parametric bootstrapping approach to determine if clinical risk factors impacted ADA frequency. Specifically, a 1,000-fold resampling procedure was performed at the cohort level, in which ADA frequencies within each clinical factor category were sampled with replacement and group medians recalculated to generate an empirical distribution of outcomes. Observed group medians (or median differences relative to a reference group) were then compared against these bootstrap distributions, and two-sided p-values were estimated as the proportion of replicates yielding medians more extreme than the observed value. This framework not only avoids assumptions of normality but also enables flexible hypothesis testing, such as assessing whether the median ADA frequency of each group significantly differs from the overall median or from a designated reference group. The factors evaluated included sampling time, dose, dosing interval, route of administration, drug mechanism of action (MOA), and co-medication. Monotonic relationships between ADA frequency and the continuous numerical factors (sampling time, dose, and dosing interval) were not assumed; therefore, these variables were converted into categorical variables by binning them into quintiles (five equal groups) for subsequent analysis. For the number of predicted T-cell epitopes, we hypothesized a monotonic, linear relationship with ADA frequency and therefore employed standard linear regression to assess this association.

To assess the combined effect of multiple variables, we performed a multivariate logistic regression. For this analysis, ADA frequency was first dichotomized into a binary outcome (low ADA vs. high ADA) using a 10% frequency cutoff. This transformation makes the analysis more robust by mitigating the influence of extreme outliers in ADA frequencies, which could otherwise disproportionately affect the regression model. The binary ADA category was then regressed against the following risk factors: drug MOA, indication, number of T-cell epitopes, dosing interval, route of administration, co-medication, dose level, and study year.

The relative importance of each risk factor was quantified by its contribution to the model’s total deviance. Deviance is a measure of model fit; a larger reduction in deviance upon adding a specific factor to the model indicates a more significant contribution of that factor to explaining the ADA outcome.

## Data availability

The entirety of the IDC DB V1 is provided in Table S4 and is openly licensed via CC BY 4.0 (creativecommons.org/licenses/by/4.0/). The IDC DB V1, future modifications and future expansions of the IDC DB will be hosted by the University at Buffalo and made accessible through the Institute for Artificial Intelligence and Data Science. All future DB versions will be memorialized and tracked and instructions for how to contribute will be provided.

## Acknowledgements

S.A. acknowledges Department of Pharmaceutical Sciences, SUNY – University at Buffalo for graduate student stipend support and his PI, Dr. Balu-iyer for guidance. Special thanks to Sathy Balu-iyer, Murali Ramanathan and the University at Buffalo for their support of this work and for hosting the IDC DB V1 and future iterations. Although they are not listed as authors on this manuscript, many individuals made important contributions to the broader IDC effort, including the initiation, design, generation, auditing, and analysis of IDC DB V1. We gratefully acknowledge: Sivan Cohen, Samuel Pine, Morten Nielsen, Gregory Steeno, Saketh Saxena, Richard Dutko, Ali Bootwala, Corey Seehus, James Brock, Quan Ho and Meghana Palaniappan.

## Supplementary information

### Supplementary Text

#### Dataset architecture, construction and quality control further explained

In addition to the Therapeutics, Sequences and Clinical Trials tables, two supporting tables were provided: Variables Explained and Controlled Language. The Variables Explained table provided definitions for each variable across the three main datasets, while the Controlled Language table standardized data entry by restricting certain fields to predefined dropdown options. For example, the field “Therapeutic Exposure Status” was limited to two controlled terms: Therapeutic Exposed and Therapeutic Naive. The purpose of this was to explicitly distinguish undosed subjects so that those receiving placebo or a different drug in the trial were not incorrectly included in the result.

Each biologic is indexed using a unique Therapeutic ID, allowing consistent traceability across therapeutic attributes, sequence-level features, and clinical trial outcomes. This relational framework enabled users to explore ADA frequency in the context of therapeutic modality, molecular characteristics, and trial-specific variables.

#### Generating aggregate datapoints at the cohort, clinical trial, therapeutic and molecule level

Supplementary Table S5 provides an aggregated ADA frequency table across four hierarchical levels: cohort, trial, therapeutic, and molecular. This table was generated by using the clinical trial dataset but with columns capturing ADA frequency estimates and corresponding identifiers for each aggregation level. At the cohort level, a unique cohort_group_id was assigned to each trial arm–exposure–drug combination, with the ADA frequency (cohort_ADA) defined as the maximum frequency of ADA-positive patients observed at any time point (max_ADA_time) within that cohort, and the corresponding number of patients (N_at_max_ADA) noted. Trial-level frequencies were calculated using a weighted average of cohort_ADA values for all cohorts within a trial_group_id (defined as trial–exposure–drug combinations), weighted by N_at_max_ADA. The total number of patients contributing to each trial-level frequency is indicated as N_of_trial_ADA. At the patient level, we report two forms of aggregate frequency: therapeutic and molecular. Therapeutic patient-level ADA frequency (PRID_ADA) was computed for each protein-exposure grouping (PR_group_id) as a weighted average of cohort-level ADA frequencies. Similarly, molecule-level ADA frequency (INN_ADA) was computed for each INN-exposure grouping (INN_group_id). The sample size used for each weighted average is reported as N_of_PRID_ADA and N_of_INN_ADA, respectively. The aggregated values did allow for comparisons across trials and molecules; however, increasing the level of aggregation obscured meaningful variation introduced by study-specific factors such as dose, regimen, or population heterogeneity.

### Supplementary Figures and Tables

**Figure S1:**
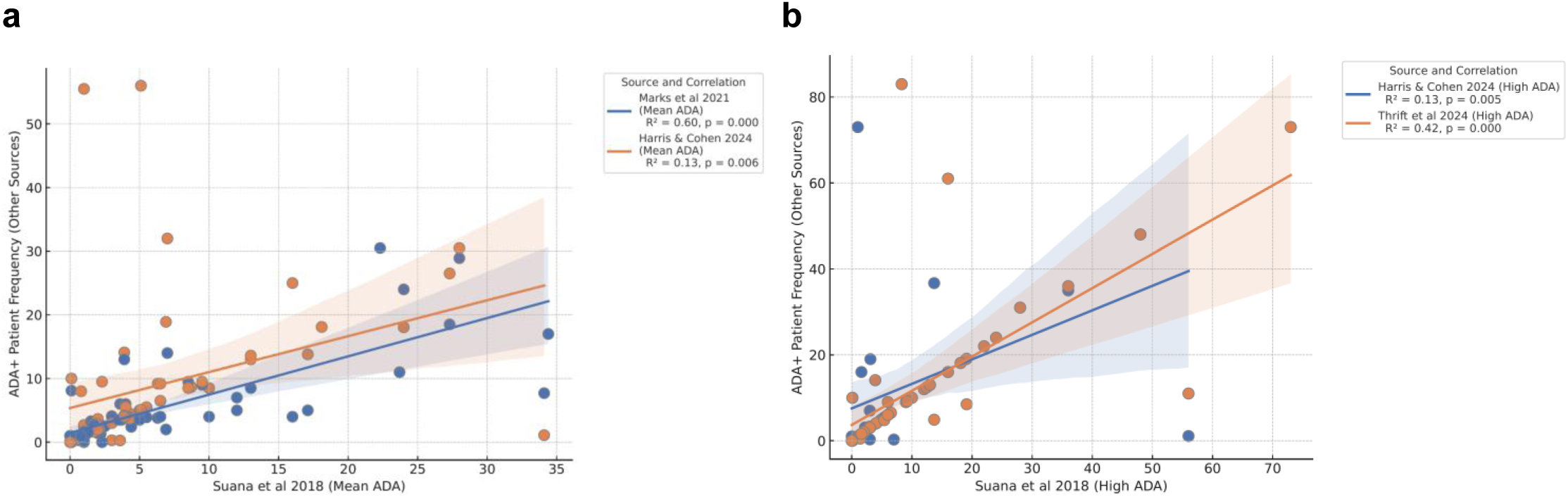
ADA frequencies reported by commonly referenced datasets. Reported ADA rates for therapeutics from commonly referenced clinical immunogenicity data table sources are compared. Suana et al 2019 = Sauna et al. Trends Biotechnol. 2018 Oct;36(10):1068-1084. doi: 10.1016/j.tibtech.2018.05.008; Marks et al 2021 = Marks et al. Bioinformatics. 2021;37(22):4041-4047. doi:10.1093/bioinformatics/btab434; Harris & Cohen 2024 = Harris CT, Cohen S. BioDrugs. 2024 2024;38(2):205-226. doi:10.1007/s40259-023-00641-2; Thrift et al 2024 = Thrift et al. Briefings in Bioinformatics. 2024;25(3)doi:10.1093/bib/bbae123. (a) Mean ADA rates reported by the indicated sources for any therapeutics shared between sources. Mean ADA frequency were either provided or calculated by taking the reported low and high ADA frequencies for each therapeutic in the data tables. (b) High ADA frequencies reported by the indicated sources for any therapeutics shared between sources. Pearson correlation best fit lines and statistics are provided.

**Table S1:**
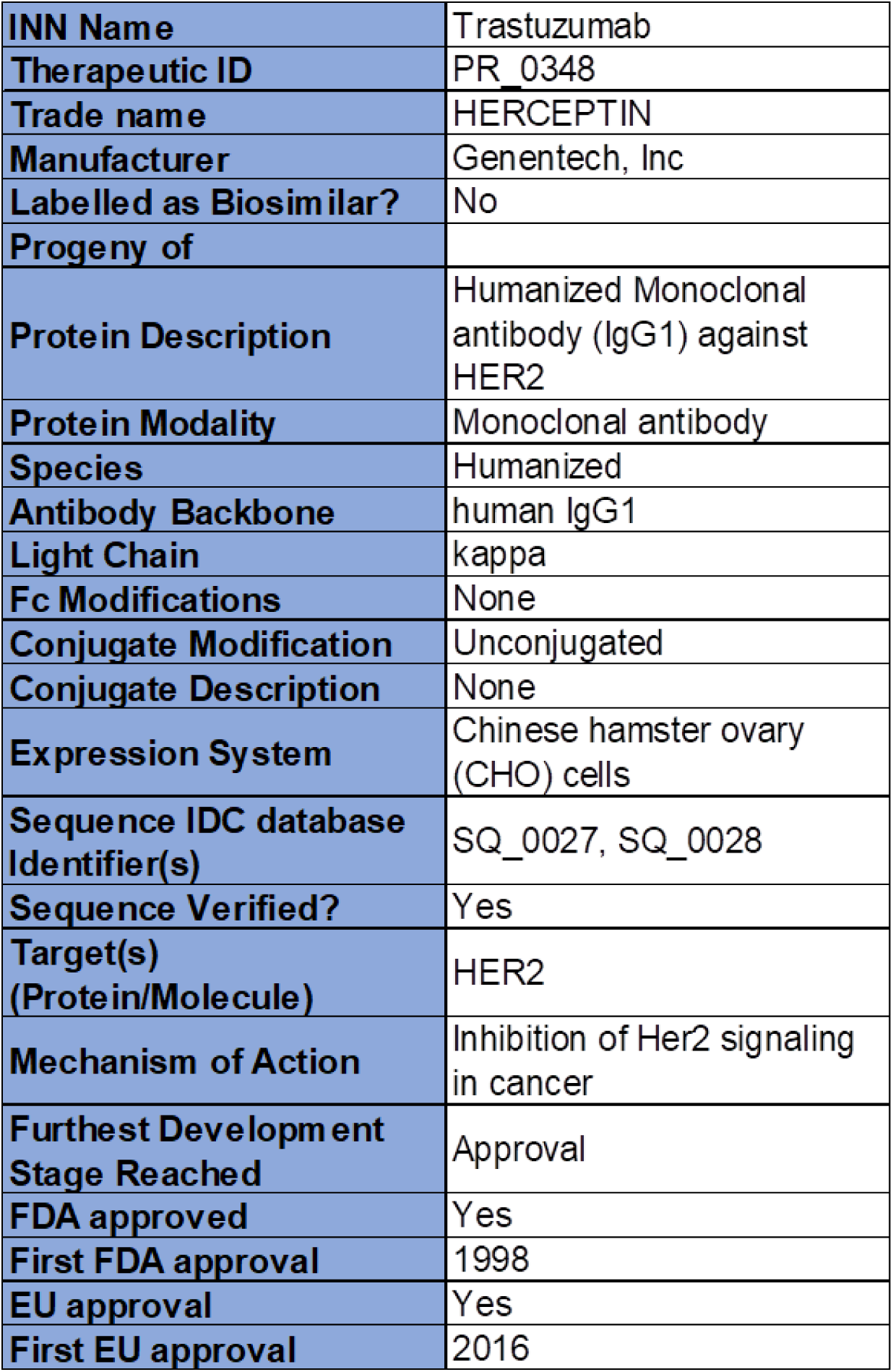
Overview of the therapeutics data table.

**Table S2:**
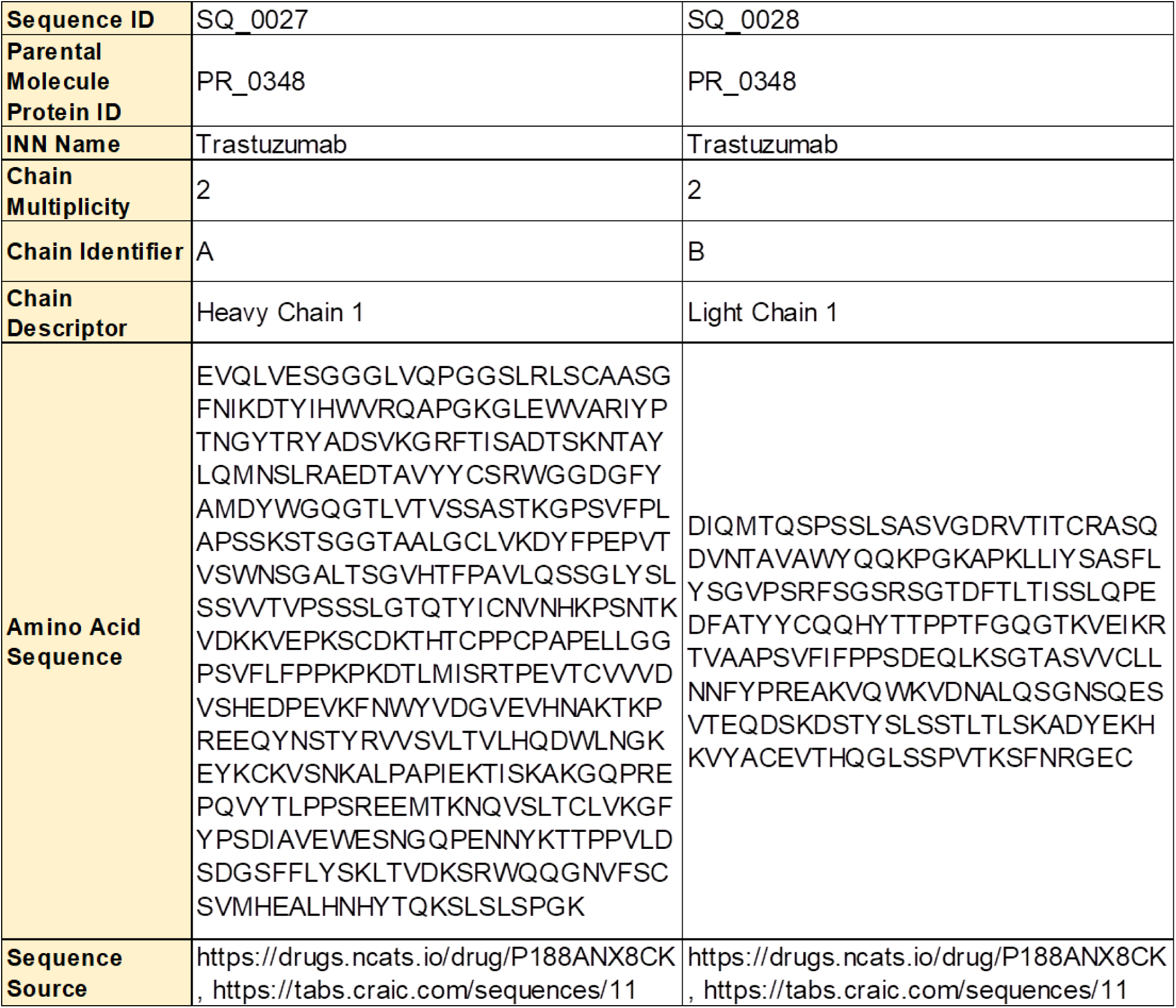
Overview of the sequences data table.

**Table S3:** Overview of the clinical trials data table.

|  |  |  |  |
| --- | --- | --- | --- |
| **Trial ID** | CT0264 | **Trial End Date** | 07-10-2014 |
| **Trial External Source** | Clinicaltrials.gov | **Therapeutic Route of Administration** | Subcutaneous |
| **External Source Identifier** | NCT01723514 | **Immunogenicity Assessment Reported Up To (Days)** | 225 |
| **Therapeutic Assessed for ADA INN Name** | Erenumab | **Number of Patients analyzed for ADA** | 6 |
| **Therapeutic Assessed for ADA Trade name** | AIMOVIG | **Number of ADA+ patients** | 0 |
| **Therapeutic Assessed for ADA ID** | PR_0139 | **Frequency of ADA+ patients** | 0 |
| **IDC Row identifier** | CT0264_A7_001 | **ADA detection assay used** | Participants who had a negative or no result at baseline and were antibody positive postbaseline. Blood samples were first tested for anti-erenumab binding antibodies, samples testing positive for binding antibodies were also tested for neutralizing antibodies. |
| **Trial Arm Identifier** | 7 | **ADA assay sensitivity or calculation** |  |
| **Immunogenicity testing** | Anti-Erenumab Antibodies | **Number of patients analyzed for ADA titers** |  |
| **Therapeutic Exposure Status** | Therapeutic Exposed | **Average ADA titers** |  |
| **Trial Arm Description** | Experimental: migraine: erenumab 140 mg | **nADAs detected** | No |
| **Trial Arm Timepoint Description** | Treatment period | **Number of patients analyzed for nADAs titers** |  |
| **Dosing Description** | Participants with migraine received 140 mg erenumab by subcutaneous injection on days 1, 29 and 57 | **Average nADA's titers** |  |
| **Therapeutic Dosing Schedule Description** | 140 mg erenumab by sc on days 1, 29 and 57 | **Number of patients analyzed for nADA** | 0 |
| **Disease Indication Category** | Neurological | **Number of patients with nADAs** | 0 |
| **Disease Indication Description** | Migraine, Healthy Subjects | **Frequency nADA+ patients reported** | 0 |
| **Co-administered drugs** | None | **Frequency of patients with hypersensitivity or injection site reactions** | Injection site pain: 0 (Frequency Threshold 5%) |
| **Patient Population** | Healthy male and female subjects, as well as male or female subjects with migraines between 18 and 55 years of age, inclusive, with no history or evidence of clinically relevant medical disorders as determined by the investigator in consultation with the Amgen physician. | **ADA interpreted to impact PK** |  |
| **Trial Start Date** | 11/14/2012 | **ADA interpreted to impact Efficacy** |  |

**Figure S2:**
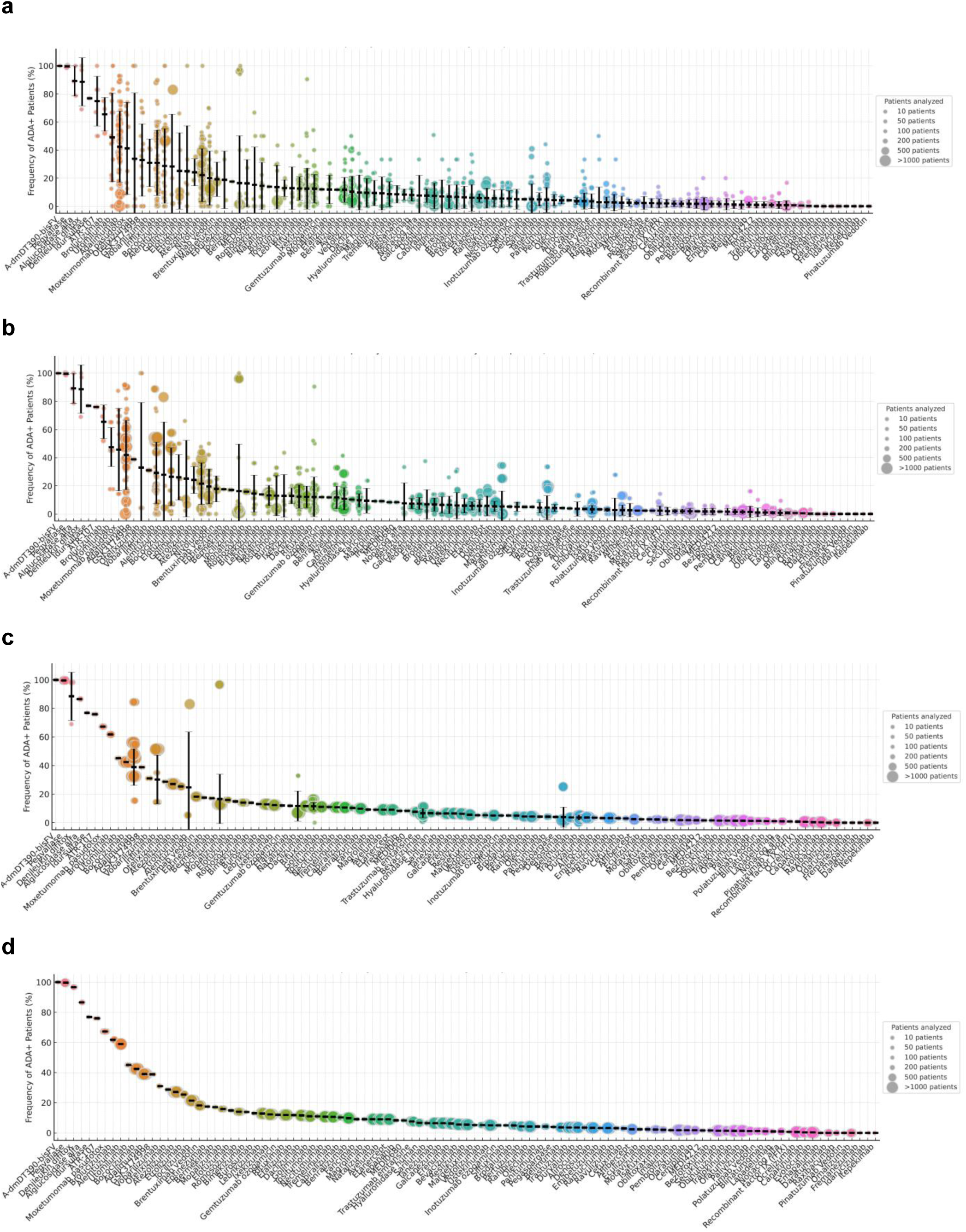
ADA Frequencies Captured in the IDC DB V1 Across All Therapeutics. (a) Cohort level ADA frequencies. Maximum observed ADA frequency is shown for each therapeutic across all clinical trial cohorts. Each data point represents a unique cohort, defined by a distinct trial-arm–exposure–drug combination (cohort_group_id, e.g., PR_0127_Therapeutic Naïve_CT0001_A1) in Table S5. ADA frequency (cohort_ADA) reflects the highest frequency of ADA-positive patients observed at any time point within the cohort. The corresponding time point is indicated by max_ADA_time, and the number of patients tested at that time point is given by N_at_max_ADA. Circle size denotes the number of patients analyzed per cohort. (b) Trial level ADA frequencies. Each point represents the trial-level ADA frequency for a therapeutic protein, derived by aggregating cohort-level data across all relevant trial arms sharing a common trial–exposure–drug grouping (trial_group_id) as seen in Table S5. ADA frequency (trial_ADA) was computed as a weighted average of maximum cohort-level ADA values, with weights corresponding to the number of patients tested at the peak time point (N_at_max_ADA) within each cohort. Circle sizes reflect the total number of patients contributing to each trial-level estimate (N_of_trial_ADA). (c) Therapeutic level ADA frequencies. Each data point represents the ADA frequency aggregated across all clinical trial cohorts associated with that therapeutic and exposure status. ADA frequency (PRID_ADA) was calculated as a weighted average of maximum cohort-level ADA frequencies (cohort_ADA), using the number of patients tested at the time of maximum ADA (N_at_max_ADA) within each cohort as weights. The protein-exposure grouping is defined by PR_group_id, and the total number of patients contributing to each value is indicated as N_of_PRID_ADA. Circle sizes reflect the patient count per therapeutic grouping. (d) Molecule level ADA frequencies. Each data point represents the ADA frequency for a therapeutic molecule, aggregated across all trials and cohorts associated with that molecule and exposure status. ADA frequency (INN_ADA) was calculated as a weighted average of maximum cohort-level ADA frequencies (cohort_ADA), using the number of ADA-tested patients at the time of maximum ADA (N_at_max_ADA) within each cohort as weights. Molecule groupings are defined by INN_group_id, and the number of patients contributing to each estimate is indicated by N_of_INN_ADA. Circle sizes correspond to the patient sample size per molecule.

**Figure S3:**
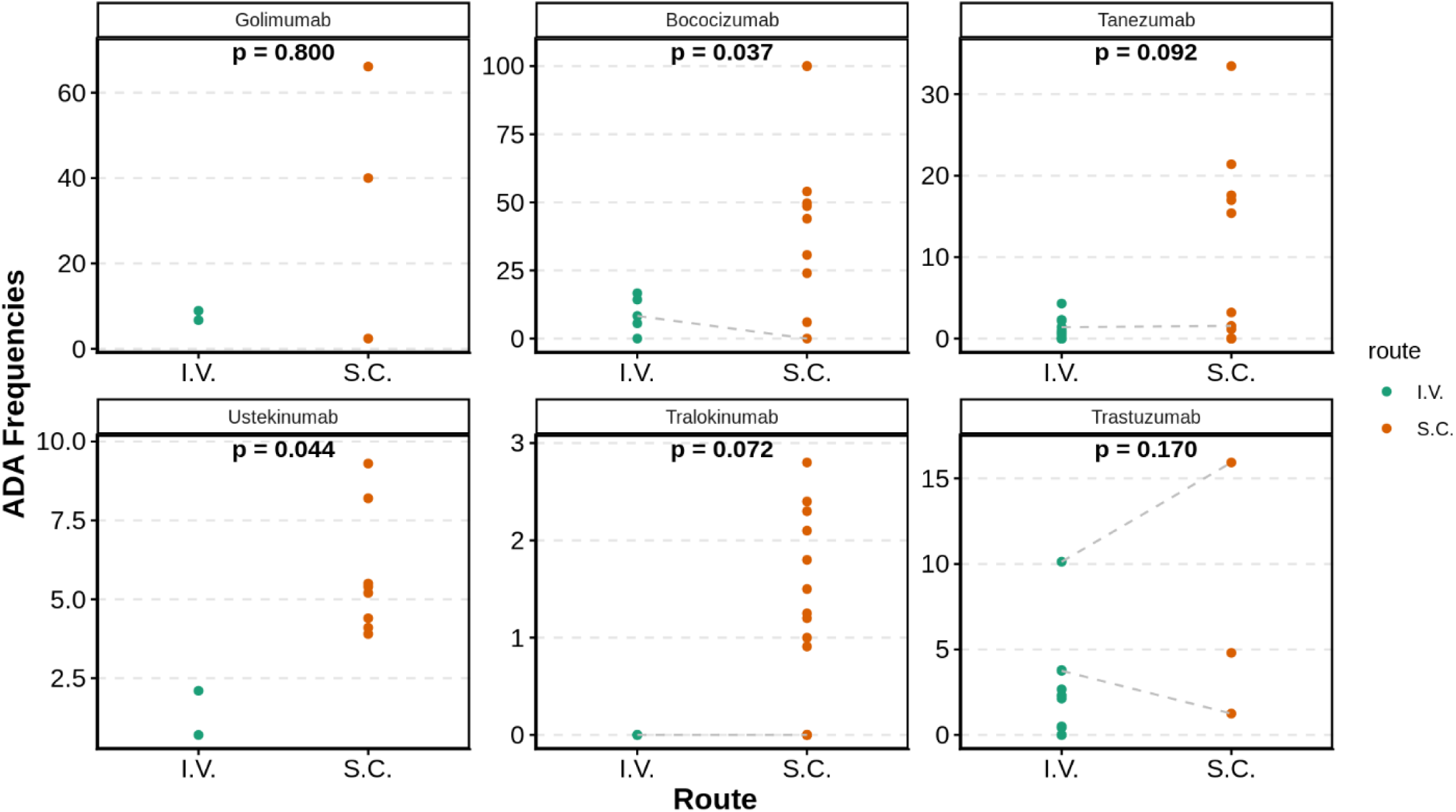
ADA frequencies comparison for SC vs IV administration.

**Figure S4:**
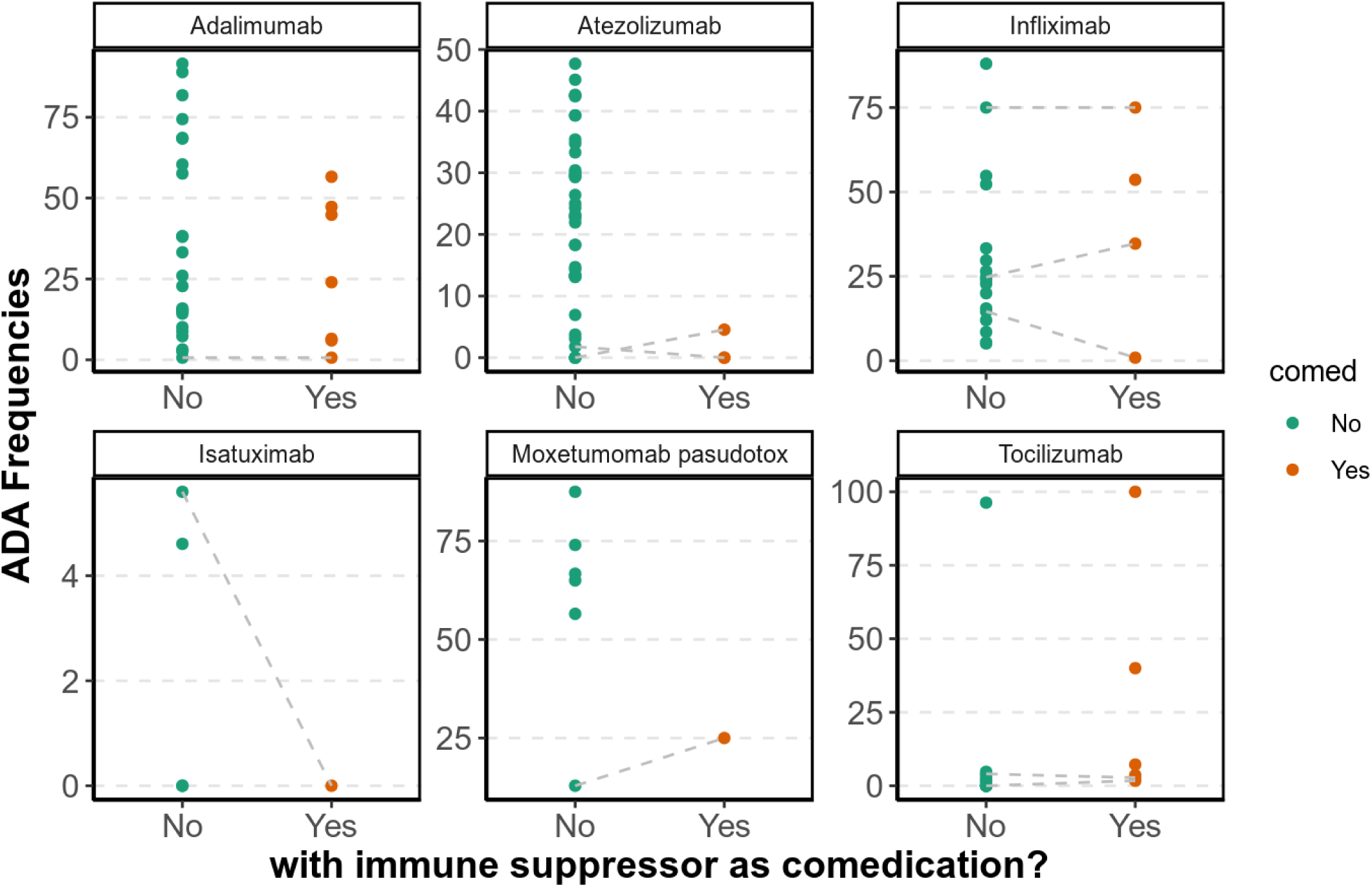
ADA frequencies for biologics evaluated in patient cohorts reported to be utilizing immune suppressive co-medications.

**Table S6:**
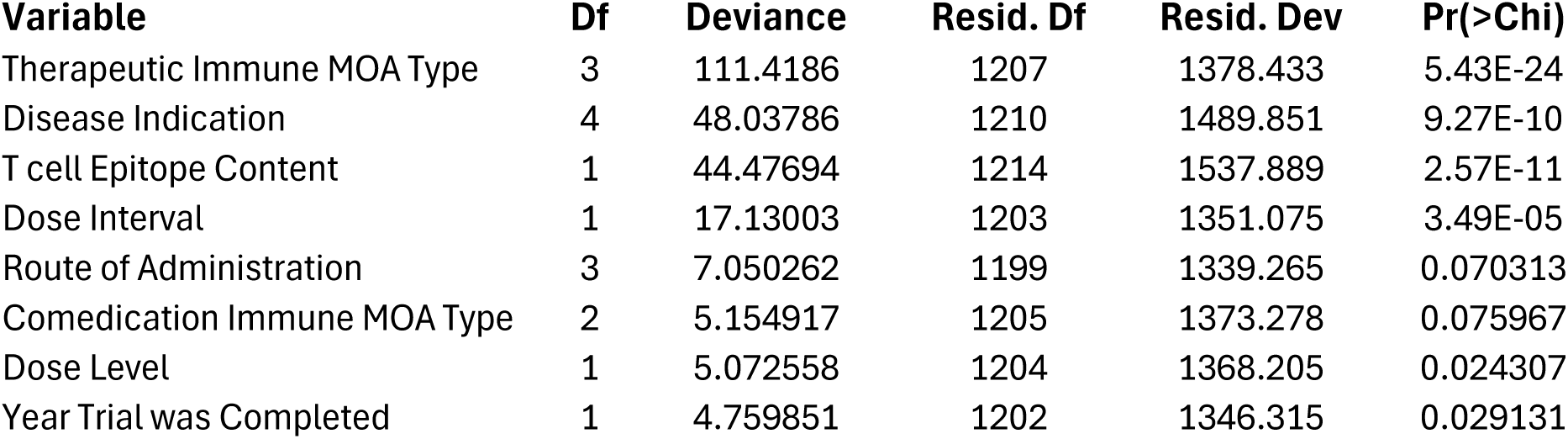
Multivariate regression results for features associated with high ADA frequency.

| Variable | Df | Deviance | Resid. Df | Resid. Dev | Pr(>Chi) |
| --- | --- | --- | --- | --- | --- |
| Therapeutic Immune MOA Type | 3 | 111.4186 | 1207 | 1378.433 | 5.43E-24 |
| Disease Indication | 4 | 48.03786 | 1210 | 1489.851 | 9.27E-10 |
| T cell Epitope Content | 1 | 44.47694 | 1214 | 1537.889 | 2.57E-11 |
| Dose Interval | 1 | 17.13003 | 1203 | 1351.075 | 3.49E-05 |
| Route of Administration | 3 | 7.050262 | 1199 | 1339.265 | 0.070313 |
| Comedication Immune MOA Type | 2 | 5.154917 | 1205 | 1373.278 | 0.075967 |
| Dose Level | 1 | 5.072558 | 1204 | 1368.205 | 0.024307 |
| Year Trial was Completed | 1 | 4.759851 | 1202 | 1346.315 | 0.029131 |

